# SPECTER-Based Semantic Triage of Biomedical Literature for Systematic Reviews in Mutational Signature Analysis

**DOI:** 10.64898/2026.07.06.736558

**Authors:** Ronie Caligagan Bituin, Ayub Bokani

## Abstract

Systematic reviews in computational biology require screening large heterogeneous bibliographic sets, especially when topics span computational methods, cancer genomics and statistical modelling. This paper presents a reproducible semantic triage pipeline that combines SPECTER scientific-document embeddings, research-question similarity, proposal-summary similarity and domain keyword coverage to rank candidate studies for systematic review screening. The pipeline was evaluated on 2,231 Covidence records, including 120 final included studies (prevalence = 5.38%), against keyword-only, TF-IDF, BM25, MiniLM, PubMedBERT and SPECTER-only baselines. SPECTER-hybrid achieved the highest average precision (AP = 0.546), recovered 50% of included studies after screening 4.48% of records, and produced an 11.16-fold enrichment over prevalence. Ablation analysis showed that semantic-keyword combinations consistently outperformed single-signal variants. These findings suggest that citation-informed hybrid ranking can support literature triage while retaining human reviewers as final decision-makers.

## I. Introduction

Systematic reviews in computational biology require reviewers to screen large volumes of biomedical literature across overlapping methodological and domain-specific areas. In mutational signature analysis, relevant studies may use terminology from cancer genomics, statistical modelling, matrix factorisation, machine learning and bioinformatics, making conventional keyword-based screening insufficient for prioritising the most relevant records.

Technology-assisted review methods can reduce manual screening burden by ranking or classifying candidate records before full reviewer assessment [9], [14], [17]. This study evaluates whether citation-informed scientific document embeddings can improve triage for a systematic review on computational methods for mutational signature analysis. The proposed SPECTER-hybrid approach combines semantic similarity to a research-question anchor, semantic similarity to a broader proposal summary and normalised domain keyword coverage. Its contribution is an empirical comparison against lexical, general embedding, biomedical embedding and SPECTER-only baselines, using average precision, precision/recall at cut-offs, NDCG, enrichment and recovery depth.

## II. Related Work

Automated and semi-automated screening has been widely studied in biomedical evidence synthesis. Earlier work explored text mining, support-vector classifiers and active learning to identify relevant studies more efficiently [14], [17], while CLEF TAR provided shared benchmarks for technology-assisted review in empirical medicine [11].

Lexical methods remain important retrieval baselines. TF-IDF ranks documents using term frequency weighted by inverse document frequency [7], while BM25 adds probabilistic term saturation and document length normalisation [6]. These methods are effective when relevant records share vocabulary with the query, but may miss studies that use different terminology for semantically related concepts.

Dense embedding models address this limitation by encoding documents into continuous vector representations. Sentence-BERT showed that transformer models can be adapted to produce cosine-similarity-preserving embeddings [16]; MiniLM provides a compact transformer model for efficient semantic similarity tasks [8]; and PubMedBERT demonstrates the value of biomedical domain pre-training [2]. However, domain-specific pre-training alone does not guarantee retrieval-optimised embeddings without similarity or ranking-oriented fine-tuning.

For scholarly retrieval, SPECTER introduced citation-informed document-level embeddings trained from scientific-paper relationships [1]. Subsequent work has examined transformer-based scientific document representations and citation-aware encoders for citation recommendation and retrieval [13], [15]. Hybrid retrieval combines sparse lexical and dense semantic signals, preserving exact domain terminology while capturing broader conceptual similarity; prior work shows that such combinations can outperform either signal alone [12].

## III. Materials and Methods

### A. Dataset and Representation

The dataset was derived from a Covidence systematic review workflow on computational methods for mutational signature analysis. The master screening file contained 2,234 bibliographic records with title, abstract, authors, year, journal and DOI. Records were labelled as included (n = 120), excluded (n = 173) or irrelevant (n = 1,938). Three ambiguous audit records were excluded, leaving 2,231 evaluation records, of which 120 were relevant (prevalence = 5.38%). These labels are reviewer-derived operational outcomes rather than independent external judgements; the pipeline is therefore evaluated as a prioritisation aid, not as an autonomous inclusion classifier.

Each record was represented by concatenating title and abstract. Transformer-based methods used raw text, while lexical methods used their internal tokenisation. Two query representations were composed: a structured research-question anchor focused on computational and statistical methods for mutational signature analysis, and a broader proposal summary covering mutation catalogues, signature decomposition and genomic data analysis.

### B. Randking Methods

Seven methods were evaluated. Keyword-only ranking counted domain terms related to mutational signatures, decomposition methods and cancer genomics. TF-IDF represented records and query text as sparse vectors with cosine similarity [7], while BM25 used the Okapi probabilistic retrieval function [6]. MiniLM used all-MiniLM-L6-v2 embeddings [8]. PubMedBERT used microsoft/BiomedNLP-PubMedBERT-base-uncased-abstract-fulltext with mean-pooled final-layer token embeddings [2], noting that this model was not retrieval fine-tuned.

The proposed SPECTER-hybrid method combines citation-informed SPECTER embeddings [1] with keyword coverage. For record i, the score was computed as:

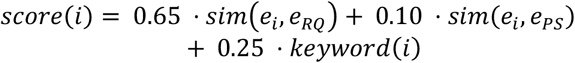

re e_i_ is the SPECTER embedding of record i, e_RQ is the research-question anchor embedding, e_PS is the proposal-summary embedding, and keyword(i) is the min-max normalised keyword coverage score. SPECTER embeddings were generated with the allenai/specter checkpoint and cosine similarity was computed using scikit-learn [5].

### C. Evaluation and Ablation

Methods were evaluated with information retrieval metrics suitable for an imbalanced dataset: average precision (AP) [4], precision at k, recall at k, NDCG at k [3], enrichment at k (P@k divided by prevalence), and recovery depth, defined as the minimum rank needed to reach 25%, 50%, 75% and 90% recall. Ablation analysis varied the weights assigned to research-question similarity, proposal-summary similarity and keyword coverage across nine configurations, including single-signal and hybrid variants. All preprocessing, ranking and evaluation steps were implemented as reproducible Python scripts; embeddings were stored as numpy arrays to allow re-evaluation without recomputation.

## VI. Materials and Methods

### A. Ranking Pefrormance

Table 1 summarises performance across seven methods on the 2,231-record dataset. SPECTER-hybrid achieved the highest AP (0.546), P@100 (0.60), Recall@100 (0.500), NDCG@100 (0.603) and Relevant@100 (60). MiniLM was the strongest single baseline, with higher P@25 and P@50 but lower AP and recovery depth than SPECTER-hybrid.

**TABLE I.** Main Ranking Performance across all Methods.

| Method | Average Precision | P@25 | P@50 | P@100 | Recall@100 | NDCG@100 | Relevant@100 |
| --- | --- | --- | --- | --- | --- | --- | --- |
| Keyword | 0.212 | 0.36 | 0.28 | 0.27 | 0.225 | 0.282 | 27 |
| TF-IDF | 0.381 | 0.52 | 0.52 | 0.52 | 0.433 | 0.513 | 52 |
| BM25 | 0.372 | 0.56 | 0.54 | 0.49 | 0.408 | 0.480 | 49 |
| MiniLM | 0.539 | 0.68 | 0.74 | 0.55 | 0.458 | 0.597 | 55 |
| PubMedBERT | 0.220 | 0.20 | 0.28 | 0.27 | 0.225 | 0.244 | 27 |
| SPECTER | 0.462 | 0.52 | 0.50 | 0.48 | 0.400 | 0.487 | 48 |
| **SPECTER-hybrid** | **0.546** | **0.60** | **0.64** | **0.60** | **0.500** | **0.603** | **60** |

**TABLE II.** Screening efficiency and enrichment at top-100 and top-200 ranked records.

| Method | Relevant@100 | Recall@100 | P@100 | Enrichment@100 | Relevant@200 | Recall@200 | P@200 | Enrichment@200 |
| --- | --- | --- | --- | --- | --- | --- | --- | --- |
| Keyword | 120 | 5.4% | 270 | 12.1% | 594 | 26.6% | 934 | 41.9% |
| TF-IDF | 59 | 2.6% | 125 | 5.6% | 422 | 18.9% | 1,002 | 44.9% |
| BM25 | 55 | 2.5% | 152 | 6.8% | 410 | 18.4% | 970 | 43.5% |
| MiniLM | 40 | 1.8% | 110 | 4.9% | 205 | 9.2% | 392 | 17.6% |
| PubMedBERT | 110 | 4.9% | 228 | 10.2% | 499 | 22.4% | 797 | 35.7% |
| SPECTER | 57 | 2.6% | 119 | 5.3% | 218 | 9.8% | 325 | 14.6% |
| **SPECTER-hybrid** | **47** | **2.1%** | **99** | **4.4%** | **187** | **8.4%** | **289** | **13.0%** |

**TABLE III.** Composition of top-100 ranked records by screening label.

| Method | Included | Excluded | Irrelevant |
| --- | --- | --- | --- |
| Keyword | 27 | 18 | 55 |
| TF-IDF | 52 | 17 | 31 |
| BM25 | 49 | 21 | 30 |
| MiniLM | 55 | 16 | 29 |
| PubMedBERT | 27 | 26 | 47 |
| SPECTER | 48 | 12 | 40 |
| **SPECTER-hybrid** | **60** | **11** | **29** |

**TABLE IV.** Ablation results for specter-hybrid weight configurations.

| Variant | w_RQ | w_Proposal | w_Keyword | AP | P@25 | P@50 | P@100 | Recall@100 | NDCG@100 | Relevant@100 |
| --- | --- | --- | --- | --- | --- | --- | --- | --- | --- | --- |
| **Full** | **0.65** | **0.10** | **0.25** | **0.546** | **0.60** | **0.64** | **0.60** | **0.500** | **0.603** | **60** |
| balanced | 0.50 | 0.20 | 0.30 | 0.543 | 0.64 | 0.64 | 0.61 | 0.508 | 0.613 | 61 |
| RQ+key | 0.75 | 0.00 | 0.25 | 0.543 | 0.60 | 0.64 | 0.59 | 0.492 | 0.595 | 59 |
| semantic-heavy | 0.75 | 0.15 | 0.10 | 0.520 | 0.60 | 0.56 | 0.55 | 0.458 | 0.570 | 55 |
| proposal | 0.00 | 1.00 | 0.00 | 0.494 | 0.68 | 0.48 | 0.53 | 0.442 | 0.542 | 53 |
| key-heavy | 0.45 | 0.10 | 0.45 | 0.475 | 0.56 | 0.50 | 0.53 | 0.442 | 0.539 | 53 |
| RQ+Proposal | 0.80 | 0.20 | 0.00 | 0.475 | 0.52 | 0.50 | 0.50 | 0.417 | 0.510 | 50 |
| RQ-only | 1.00 | 0.00 | 0.00 | 0.462 | 0.52 | 0.50 | 0.48 | 0.400 | 0.487 | 48 |
| key-only | 0.00 | 0.00 | 1.00 | 0.211 | 0.36 | 0.28 | 0.26 | 0.217 | 0.275 | 26 |

TF-IDF and BM25 outperformed keyword-only ranking, confirming the value of term weighting beyond simple term counts. SPECTER alone exceeded both lexical baselines, while PubMedBERT performed close to keyword-only ranking, consistent with the use of mean-pooled encoder representations without retrieval-oriented fine-tuning.

**Fig. 1.**
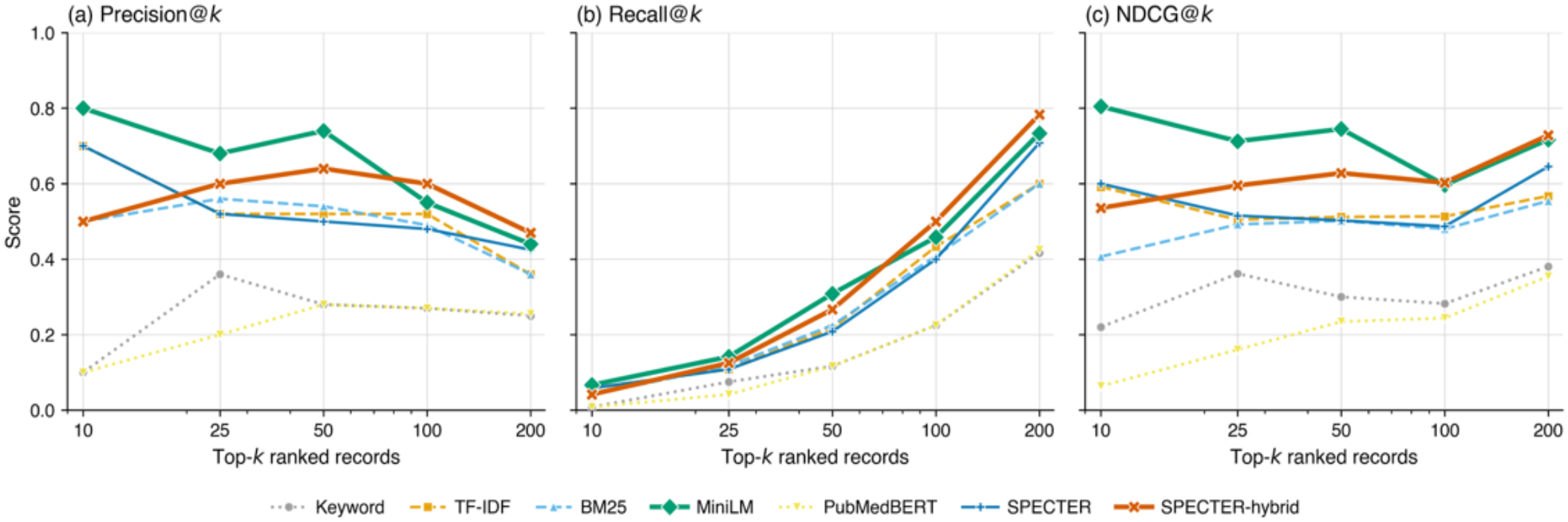
Precision@k (a), Recall@k (b), and NDCG@k (c) across screening cut-offs k ∈{10, 25, 50, 100, 200} for all seven ranking methods.

### B. Ranking Pefrormance

SPECTER-hybrid concentrated relevant studies most strongly at the top of the ranked list. At the top 100 records, it recovered 60 of 120 final included studies after screening 4.48% of the evaluation set, corresponding to 11.16-fold enrichment over prevalence. At the top 200 records, it recovered 94 studies (recall = 0.783).

Recovery depth confirmed the workload advantage. SPECTER-hybrid required the fewest records to reach every threshold: 50% recall at rank 99, 75% at rank 187 and 90% at rank 289. MiniLM required ranks 110, 205 and 392 for the same thresholds, while lexical methods fell substantially behind at higher recall levels.

### C. Top-100 Composition by Method

To examine whether the ranking methods merely retrieved reviewed-but-rejected studies or genuinely concentrated final included records, the top 100 records for each method were grouped by original screening label. SPECTER-hybrid produced the highest density of included studies in the top 100 while also yielding the lowest count of excluded records, suggesting that the hybrid score improved both relevance concentration and screening usefulness.

**Fig. 2.**
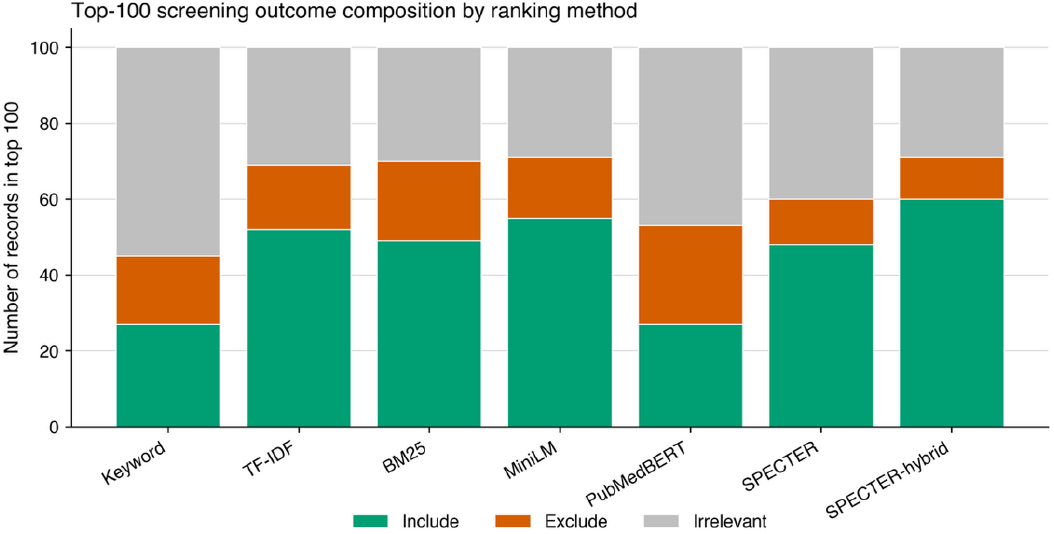
Stacked bar chart showing the composition of the top-100 ranked records by screening outcome label (included, excluded, irrelevant) for each ranking method.

SPECTER-hybrid ranked 60 included studies, 11 excluded studies and 29 irrelevant records in the top 100. SPECTER alone ranked 40 irrelevant records in the top 100, more than MiniLM and SPECTER-hybrid, indicating that keyword coverage helps suppress topically distant records that may share surface-level embedding similarity with the query.

### D. Ablation Analysis

**Fig. 3.**
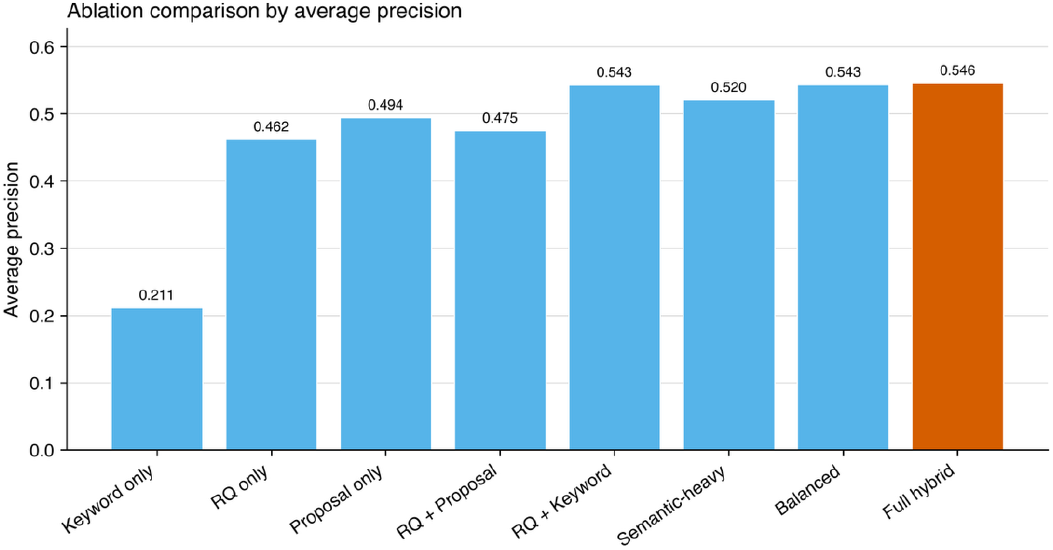
Average precision for each hybrid weight configuration in the ablation study. The full hybrid configuration (orange) achieved the highest average precision (AP = 0.546).

Ablation results showed that hybrid combinations consistently outperformed single-signal variants. The full configuration achieved the highest AP (0.546), while the balanced variant achieved the highest NDCG@100 (0.613) and retrieved 61 relevant records in the top 100. Reducing keyword weight to 0.10 lowered AP to 0.520, whereas over-weighting keywords to 0.45 lowered AP to 0.475. Thus, domain keyword coverage is complementary to semantic similarity, but works best at moderate weight.

## V. Discussion

The results demonstrate that combining SPECTER scientific-document embeddings with domain keyword coverage improves literature prioritisation compared with either signal alone. SPECTER captures broad semantic and topical similarity based on citation context, whereas keyword coverage anchors the score to the vocabulary of the specific review question. This pattern is consistent with hybrid retrieval literature showing that dense and sparse signals are complementary [12].

MiniLM was a strong competitor, achieving AP = 0.539 and outperforming SPECTER-hybrid at early precision cut-offs. One plausible explanation is that MiniLM is contrastively trained for short text similarity, while SPECTER is designed for document-level scientific relatedness. However, SPECTER-hybrid performed better at AP, P@100, NDCG@100 and recovery depth, suggesting that keyword coverage helps maintain ranking quality deeper in the screening queue. PubMedBERT performed poorly despite biomedical pre-training [2], supporting the view that retrieval-oriented fine-tuning, rather than domain pre-training alone, is critical for cosine-similarity retrieval [16].

Practically, the pipeline is intended to support rather than replace human reviewers. Reaching 75% recall after screening 8.4% of records and 90% recall after 13.0% indicates meaningful workload reduction, while all final inclusion decisions remain with reviewers applying eligibility criteria. This aligns with TAR evaluation frameworks that assess the review effort required to reach target recall levels [10].

This study contributes a domain-specific evaluation of SPECTER-based hybrid ranking for computational biology systematic reviews, where relevant studies span computational methods, cancer genomics and statistical modelling. Unlike purely lexical triage, the hybrid approach can prioritise records that are conceptually relevant even when terminology varies, while still rewarding domain-specific vocabulary.

## VI. Limitation

Several limitations should be noted. The evaluation used a single systematic review dataset, so generalisation to other topics, eligibility criteria or reviewer teams requires further validation. The labels are reviewer-derived workflow outcomes rather than independent ground truth, and the exploratory notebook also supported screening, introducing possible circularity. PubMedBERT was not retrieval fine-tuned, so stronger biomedical sentence-transformer baselines may perform better. The keyword component depends on a manually curated domain list, and the hybrid weights were selected empirically on the same dataset rather than on a held-out review. Finally, the pipeline has not yet been evaluated on shared benchmarks such as CLEF TAR [11].

## VII. Conclusion

This paper presented a SPECTER-based hybrid semantic triage pipeline for prioritising biomedical literature in systematic review screening. On 2,231 records from a mutational signature analysis review, the method achieved the highest average precision among seven methods and recovered 50% of final included studies after screening fewer than 5% of records. Ablation analysis showed that the advantage came from combining semantic and lexical signals: neither SPECTER embeddings alone nor keyword coverage alone matched the hybrid formulation. These results support hybrid semantic-keyword ranking as a practical, reproducible triage aid for computational biology reviews, with future work focused on retrieval-fine-tuned biomedical embeddings, external benchmarks and prospective validation.

